# Canagliflozin reprograms the aging hippocampus in genetically diverse UM-HET3 mice and attenuates Alzheimer’s-like pathology

**DOI:** 10.1101/2025.06.10.658747

**Authors:** Hashan Jayarathne, Dulmalika Herath Manchanayake, Nina Chimienti, Omar Kadri, Katherine Gurdziel, Seongho Kim, Hyejeong Jang, Brett C. Ginsburg, Richard A. Miller, Shoshana Yakar, Marianna Sadagurski

## Abstract

Aging is the strongest risk factor for cognitive decline and Alzheimer’s disease (AD), yet the mechanisms underlying brain aging and their modulation by pharmacological interventions remain poorly defined. The hippocampus, essential for learning and memory, is particularly vulnerable to metabolic stress and inflammation. Canagliflozin (Cana), an FDA-approved sodium-glucose co-transporter 2 inhibitor (SGLT2i) for type 2 diabetes, extends lifespan in male but not female mice, but its impact on brain aging is unknown. Here, we used a multi-omics strategy integrating transcriptomics, proteomics, and metabolomics to investigate how chronic Cana treatment reprograms brain aging in genetically diverse UM-HET3 mice. In males, Cana induced mitochondrial function, insulin and cGMP–PKG signaling, and suppressed neuroinflammatory networks across all molecular layers, resulting in improved hippocampal-dependent learning and memory. In females, transcriptional activation of neuroprotective pathways did not translate to protein or metabolite-level changes and failed to rescue cognition. In the 5xFAD AD model, Cana reduced amyloid plaque burden, microgliosis, and memory deficits in males only, despite comparable peripheral glucose improvements in both sexes. Our study reveals sex-specific remodeling of hippocampal aging by a clinically available SGLT2i, with implications for AD pathology and lifespan extension, and highlights Cana’s potential to combat brain aging and AD through sex-specific mechanisms.

## Introduction

Aging is a significant risk factor for cognitive decline and neurodegenerative disorders, including Alzheimer’s disease (AD)(1). Patients with AD exhibit distinct pathological changes, such as filamentous inclusions of the microtubule-associated protein tau, extracellular deposits of amyloid-beta (Aβ) in senile plaques, pronounced neuroinflammation, and loss of neuronal cells(2). Although considerable efforts have been made to address amyloid and tau pathologies, an increasing body of evidence now suggests that metabolic dysfunction and neuroinflammation play crucial roles in the initiation and progression of the disease(3, 4). Supporting this, individuals with insulin resistance, obesity, and metabolic disorders have a higher risk of developing AD(5). Within the central nervous system (CNS), the hippocampus is particularly susceptible to metabolic changes due to its high energy demands for memory and learning processes, and hippocampal changes have been linked to cognitive impairment, dementia, and AD(6). While there are currently no effective treatments for AD, it has been suggested that drugs that improve metabolic health, reduce age-related neuroinflammation, and enhance both healthspan and lifespan might be repurposed to treat neurodegenerative diseases like AD(1).

The Interventions Testing Program (ITP) has identified several treatments that successfully extend lifespan and improve metabolic health in the genetically diverse UM-HET3 mice, including the sodium-glucose co-transporter 2 inhibitor (SGLT2i) Canagliflozin (7–10). Canagliflozin (Cana), an FDA-approved drug, is commonly used to treat type 2 diabetes (T2D)(11). SGLT2i act by blocking glucose reabsorption in the renal proximal tubules, leading to increased urinary glucose excretion and improved glycemic control through an insulin-independent mechanism(11). Randomized clinical studies indicated that SGLT2i exhibit numerous pleiotropic effects and benefits, including cardiovascular and all-cause mortality reduction, independent of their impact on diabetes(12).

In mice, Cana extended lifespan by 14% and robustly retarded age-related lesions in all tissues of male but not female mice (10, 13). We showed that long-term (∼20 months) Cana treatment significantly reduced neuroinflammation in the hippocampus and hypothalamus, improved metabolic health, neuromuscular function, locomotor activity, and anxiety-like behavior in 30-month-old male mice(14). This phenotype indicated that Cana exerts broad neuroprotective effects on brain regions critical for cognitive function and metabolic regulation. However, the evidence that Cana benefits males far more than females emphasize a crucial gap in our understanding of how therapeutic interventions perform across sexes. These findings support a strong rationale for repurposing Cana to prevent or delay age-associated neurodegenerative diseases.

The molecular mechanisms of Cana’s effects on the hippocampus and its impact on cognition and central metabolism remain largely unexplored. We employed a multi-omics strategy, integrating transcriptomics, proteomics, and metabolomics, to obtain a system-level understanding of the complex biological changes induced by Cana treatment in the aged hippocampus. Multi-omics analyses offer the advantage of providing a more detailed understanding of disease pathogenesis, and they have been widely applied to brain aging and neurodegenerative studies (15). For example, multi-omics analysis (metabolomics and transcriptomics) revealed the presence of neuroinflammation, activated glial signaling, dysregulated synaptic signaling, and impaired metabolism in the hippocampus of mice between 6, 12, and 24 months of age (16). Our integrative approach revealed that long-term Cana treatment reprograms hippocampal molecular pathways in a sex-specific manner, supporting improved cognitive function and reduced neuroinflammation in aged males.

To validate our findings, we studied AD progression in 5xFAD mice treated with Cana. The 5xFAD mice overexpress the human amyloid precursor protein (APP) containing three familial Alzheimer’s disease (FAD) mutations, and human presenilin-1 (PSEN1) with two additional FAD mutations. These mice exhibit hallmark features of AD, including cognitive deficits, amyloid plaque deposition, sex-specific differences, and neuroinflammatory changes from 4 to 12 months of age, recapitulating many aspects of human AD pathology (17, 18). Data from our study establishes a link between glucose metabolism and cognitive function in aging, demonstrating that Cana, which targets peripheral and central glucose metabolism, can delay cognitive decline and neuropathological features associated with AD. Our results position Cana as a promising candidate for therapeutic repurposing in the context of age-related neurodegeneration.

## Results

### Cana treatment improves cognitive function in middle-aged male mice

To assess the impact of Cana on cognitive and neuromotor function, UM-HET3 mice were treated from 7 months of age and subjected to behavioral testing at 12-14 months (**Figure 1A**). Cana-treated male but not female mice spent significantly more time in the center of the open field (p < 0.05), indicating reduced anxiety-like behavior (**Figure 1B**). Notably, the total distance traveled did not differ between groups, suggesting that locomotor activity was not yet affected at this age (**Figure 1C**). To further assess the impact of Cana on cognitive function, we evaluated spatial learning and memory by measuring the latency to find the target hole over 4 days of training in the Barnes maze (**Figure 1D**). During the acquisition phase (training days 1–4), Cana-treated males spent significantly less time finding the target hole than controls (**Figure 1E**), indicating accelerated learning. This effect was only significant on day 4 (p < 0.05) for Cana-treated females (**Figure 1F**). Spatial working memory (short-term and long-term memory, STM and LTM) tests were performed on days 5 and 12 to assess reference memory for the previously learned target hole. On both days, Cana-treated males located the previously learned target hole significantly faster than control males (p < 0.01) and spent more time in the target quadrant (p < 0.05), indicating enhanced memory retention (**Figure 1G**). These outcomes show a significant effect of sex (p < 0.01) and sex × Cana diet interaction (p < 0.05). Together, these results demonstrate that Cana improves hippocampal-dependent learning and memory and reduces anxiety-like behavior, particularly in middle-aged males.

**Figure 1:**
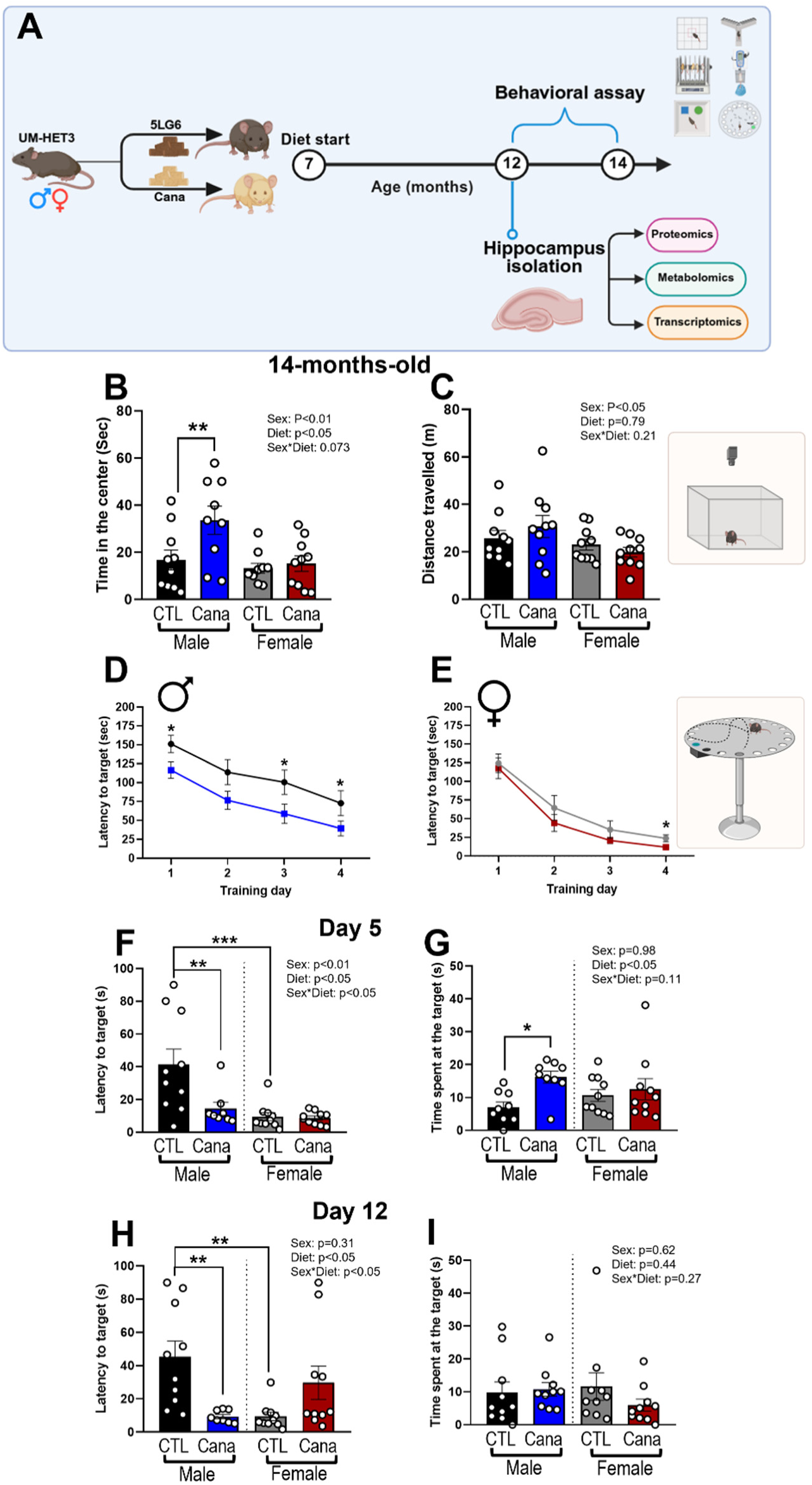
Cana improves cognitive function in middle-aged male mice. A) Schematic diagram of experimental design. 7-month-old UM-HET3 male and female mice were randomly assigned to control or Cana diet. Behavioral testing was conducted at 14 months of age. B) Time spent in the center (seconds); C) Distance travelled (meters) during the open field test. Latency to target hole (seconds) during training day 1-4 D) Males; E) Females. F) Latency to target hole (seconds); G) Time spent at the target hole on the 5^th^ day for short-term memory test. H) Latency to target hole (seconds); I) Time spent at the target hole on the 12^th^ day for long-term memory test. Error bars show SEM for n = 8–10 mice/group. Data was analyzed using Two-factor ANOVA and further analyzed with the Newman–Keuls post hoc test (*p < 0.05, **p<0.01, ***p<0.001).

### Hippocampal proteomics analysis in mice treated with Cana

Cana levels in the hippocampus were measured using high-performance liquid chromatography/mass spectrometry (19). Consistent with our previous studies (14), Cana levels in the hippocampus were significantly higher in females than in males (**Figure 2A**). Cognitive function and behavior are controlled by the hippocampus (20). We took an unbiased hippocampal proteomics approach to understand the molecular mechanisms underlying improved behavior in Cana-treated mice. We identified 129 down-regulated and 157 up-regulated proteins in Cana-treated males and 56 down-regulated and 97 up-regulated proteins in Cana-treated females (**Figure 2B and C**). Principal Component Analysis (PCA) revealed distinct responses to Cana in each group (**Suppl Figure 1A and B**). Gene Ontology (GO) analyses of differentially expressed proteins revealed enrichment in Cognitive function and Metabolism pathways in Cana-treated male mice. Specifically, the ‘Negative regulation of beta-amyloid formation’ pathway was the most enriched in Cana-treated males, showing a dramatic 40-fold enhancement compared to controls (**Figure 2D**). Top upregulated proteins included PIN1, BIN1, RTN3, GGA3, and SORL (**Figure 2F and I**), known to inhibit amyloidogenic APP processing and reduce Aβ accumulation (21–24). Additional altered proteins involved in metabolic pathways, including catabolic processes, lipid metabolism, carbohydrate metabolism, and fatty acid metabolic processes, are shown in the heatmap (**Figure 2G and J**). Cana-treated females exhibited enrichment of pathways primarily associated with metabolic function, including ‘fatty acid biosynthesis’, ‘ATP synthesis’, ‘fatty acid metabolism’, and ‘estrogen biosynthetic process’ (**Figure 2E**). This enrichment was driven by the upregulation of key metabolic proteins such as ATP5E, ATP5I, MECR, NDUDBB, S27A4, and ASAH1 (**Figure 2H and K**), suggesting enhanced metabolic resilience induced by Cana treatment in female mice (25, 26).

**Figure 2:**
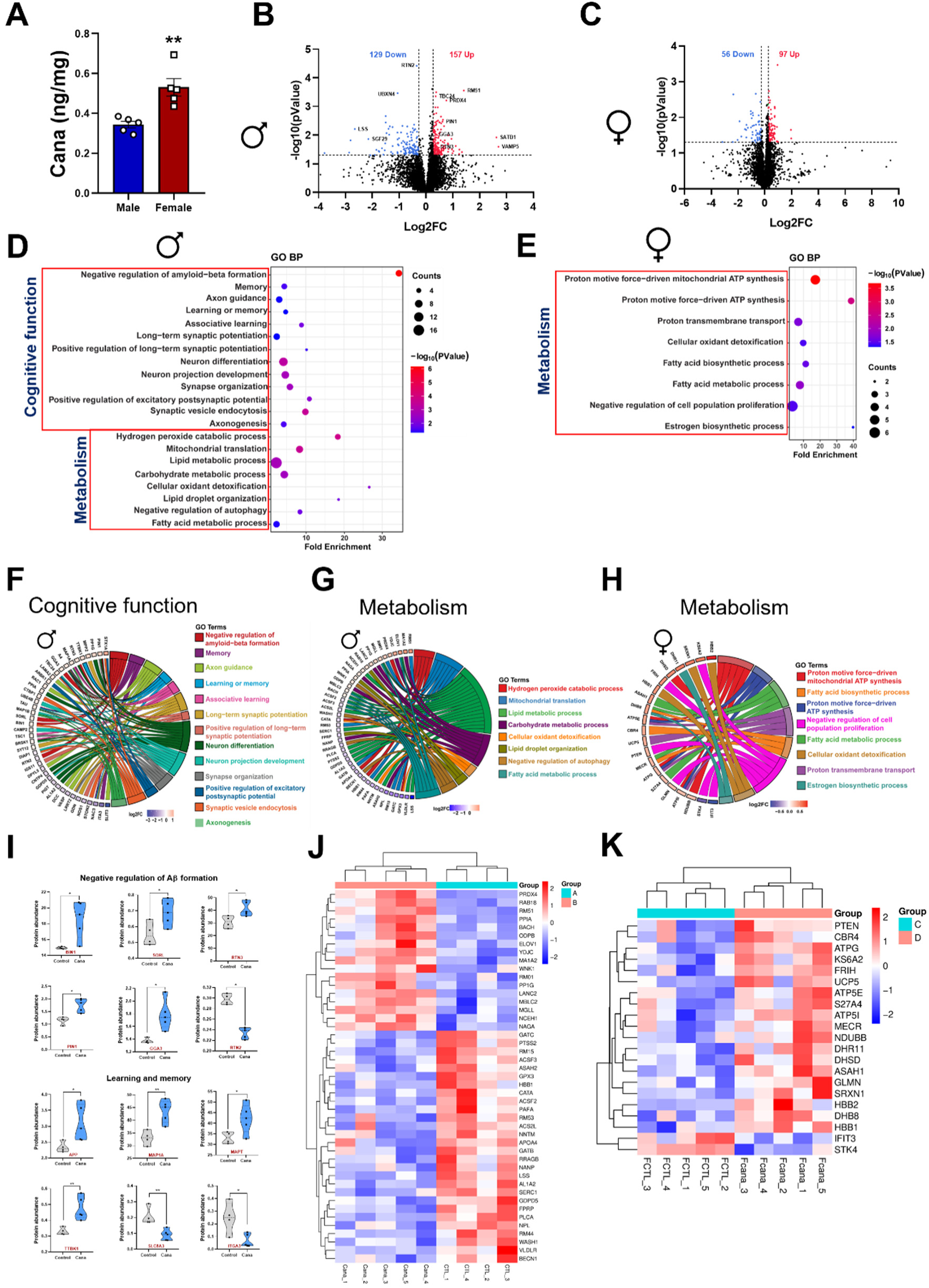
Cana modulates the hippocampus proteome in 12-month-old mice. **(A)** Bar graph showing the Cana levels in the hippocampus. Volcano plot showing differentially expressed proteins in B) Males; C) Females. Gene ontology (GO) analysis showing biological processes was analyzed using differentially abundant proteins in D) Males, and E) Females. GO chord showing the directionality of the proteins relevant to each pathway, broadly categorized as Cognitive function, and Metabolism-related pathways, F) and G) Males, and H) Females. I) Violin plot showing proteins in highly enriched ‘negative regulation of amyloid beta formation’ pathway and ‘Learning and memory’ proteins in males. Heat maps show all the proteins categorized under Metabolism in J) Males, and K) Females. n=5 mice/group

### Hippocampal metabolomic analysis in mice treated with Cana

The sex-dependent enriched metabolic pathways seen by proteomics prompt us to assess the metabolome of the hippocampus. We performed untargeted hippocampus metabolomics of 12-month-old mice. PCA demonstrated distinct clustering between male and female treatment groups, indicating different responses (**Suppl. Figure 1C and D**). We identified 103 differentially regulated metabolites in Cana males and 142 in Cana females compared to controls. Both sexes exhibited an increased abundance of amino acid metabolites and decreased carbohydrate metabolites (**Figure 3A and D**), suggesting a metabolic shift toward amino acid utilization in the hippocampus. Importantly, in Cana males, the top-ranked upregulated metabolites, such as phosphatidylethanolamine, n-arachidonoyl taurine, palmitoylethanolamide, and docosatrienoic acid (**Figure 3C**), were all related to anti-inflammatory signaling and neuroprotection(27–30). Additionally, metabolite set enrichment analysis (MSEA) in males revealed enrichment in pyrimidine and glutathione metabolism (**Figure 3E**), critical for neuronal repair and oxidative resilience(31, 32). In Cana-treated females, most upregulated metabolites were related to lipid metabolism and remodeling, such as C24:1 sphingomyelin, gingerglycolipid C, and 1-stearoyl-2-myristoyl-sn-glycero-3-phosphocholine (**Figure 3D**). MSEA in females indicated activation of sphingolipid pathways and amino sugar metabolism (**Figure 3F**). Only two downregulated metabolites, erythrose and 1-(9Z-octadecenoyl)-2-(9Z,12Z-octadecadienoyl)-sn-glycero-3-phosphocholine (PC(18:1/18:2)), were shared between Cana-treated male and female mice, indicating sex-independent metabolic shifts induced by Cana. A male sex-specific, adrenosterone, and leu-Gly-His with known anti-neuroinflammatory properties(33), were upregulated in males but downregulated in females, indicating a sex-specific impact of Cana on hippocampal metabolism (**Figure 3I**).

**Figure 3:**
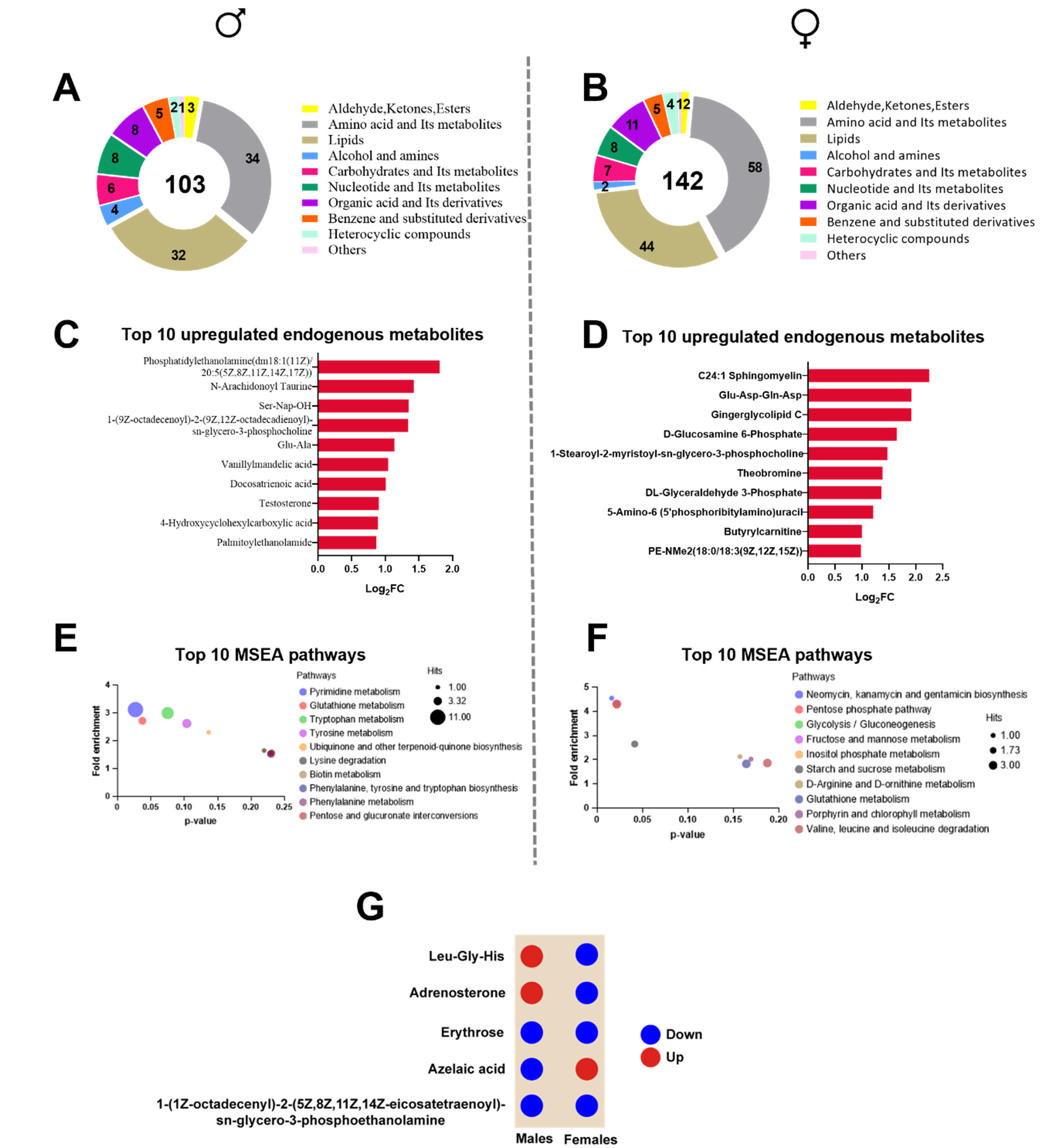
Hippocampus metabolome analysis in Cana-fed mice. Differentially expressed metabolites and their categories in the Cana-fed A) Male; B) Female mice detected by GC-MS. Top 10 upregulated metabolites in C) Males; D) Females. Top 10 pathways in metabolite set enrichment analysis (MSEA) in Cana fed E) Male mice; F) Females. G) Shared metabolites between males and females and their expression direction. n =4/5 mice/group.

### Hippocampal transcriptomics analysis in mice treated with Cana

To understand Cana’s effects on transcriptional remodeling, we performed bulk RNA sequencing of hippocampal tissue from mice treated with Cana at 12 and 25 months. At 12 months, differential gene expression analysis revealed 403 upregulated and 469 downregulated genes in Cana-treated males, and 631 upregulated and 834 downregulated genes in females (**Figure 4A and D**). Of these, 123 genes were shared between sexes, indicating that there are both shared and sex-specific transcriptional responses. KEGG and GO pathway analysis of the shared genes revealed common enrichment in behavioral and gene regulatory processes, including ‘transcriptional control’, ‘MAPK cascade regulation’, and ‘drinking behavior (**Figure 4H**), likely reflecting a direct effect of SGLT2i on water consumption(34). PCA plots are shown in **Suppl. Figure 1E and F**. In males, transcriptomic remodeling was characterized by strong enrichment in KEGG pathways related to ‘oxidative phosphorylation’, ‘circadian rhythm’, and neurodegenerative diseases, including ‘Alzheimer’s disease’. Cana downregulated key genes of inflammatory and insulin-related signaling (*Socs1*, *Irs2*, *Irs4*, *Rps6ka4*, *Jund*, *Akt1s1*, *Map2k2*, and *Bace1*), while upregulated genes were associated with metabolic regulation and neuroprotection (*Irs1*, *Phkg1*, *Flot1*, *Casp9*, *Csnk2b*, and *Wnt11*) (**Figure 4B and C and suppl. Figure 2A**). Cana-treated females showed enriched insulin, MAPK, mTOR, glutamatergic, GABAergic, and longevity-related signaling pathways (**Figure 4E**). This enrichment was largely driven by the downregulation of critical genes involved in insulin signaling (*Akt2*, *Irs3*, *Pdk4*), synaptic function (*Grm3*, *Gabrd*, *Gad2*), and adaptive plasticity (*Arc*, *Calb1*), suggesting a sex-specific transcriptional response in females (**Figure 4F and G**). At 25 months, Cana-treated males maintained the transcriptional changes observed at 12 months. PCA plots are shown in **Suppl. Figure 1G and H**. Specifically, we observed enrichment in ‘nervous system development’ and ‘neuronal differentiation’, ‘synaptic plasticity’, ‘insulin signaling pathway’, and ‘innate immune response’, with suppression of neuroinflammatory genes such as *Nod2*, *Il34*, and *Parp14* (**Figure 4I and J, and Suppl. Figure 2B and C).** GO analysis of 25-month-old Cana-treated females revealed enrichment in cognition and synaptic activity processes, including ‘locomotor behavior’, ‘associative learning’, ‘axon guidance’, ‘long-term memory’, and ‘synaptic potentiation’. These pathways were largely driven by downregulating the key genes such as *Egr1*, *Arc*, *Calb1*, *Fos*, *Abcc8*, *Prkacb*, *Itpr1*, and *Camk4*. Interestingly, pathways related to ‘Insulin receptor’ and ‘MAPK signaling’ pathways were also enriched, with downregulation of *Akt2* and *Foxo3*, and upregulation of *Irs1* genes. We detected a cluster of up- and downregulated immune and inflammatory genes in aged Cana-treated females (**Figure 4K and L, and Suppl. Figure 2D)**. Across both age groups, Cana males exhibited 38 shared genes, including upregulated mitochondrial and metabolic genes such as *Creb1*, *Irs1, Irs2*, *Smn1*, *Cd84*, *Ccr2*, *mt-Nd2*, *mt-Nd4*, *mt-Nd5*, *P2ry13*, and *Slc10a3*, *Slc46a3*, as well as downregulated pro-senescent and inflammatory genes like *Glo1*, *Pm20d1*, and *Insig1* **(Figure 4M)**. In contrast, among the shared genes for females at both age groups (12 and 25 months), we observed the consistent upregulation of chromatin modifiers and DNA damage genes (*Bend3*, *Zbtb16* and *Parp9*). Meanwhile *Akt2*, *Calb1*, *Egr1*, *Egr4*, *Hhex*, and *Lrrk2* remained downregulated, suggesting a reduced plasticity and signaling (**Figure 4N**).

**Figure 4:**
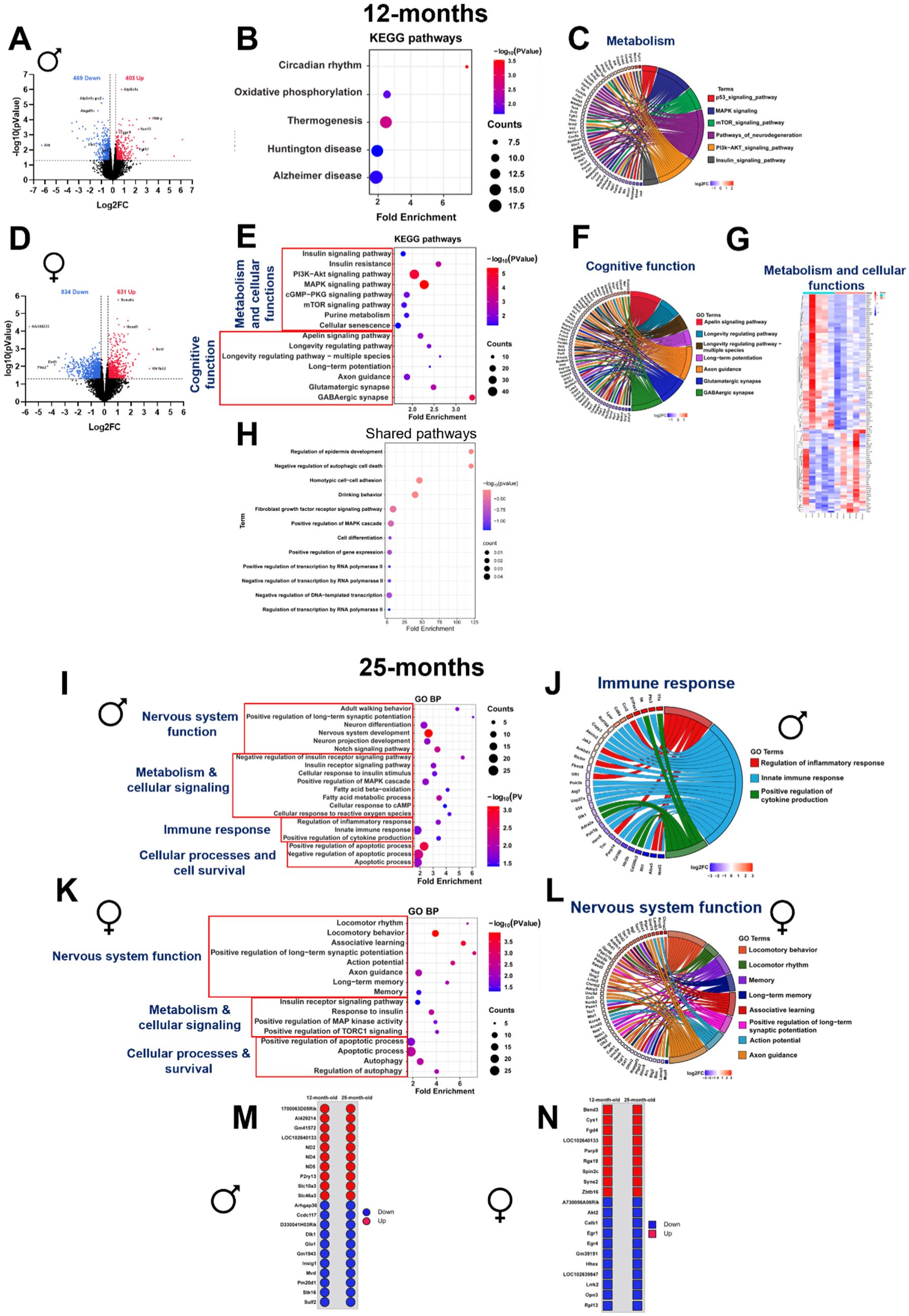
Sex-specific transcriptomic changes induced by Cana in 12 and 25-month-old mice. Volcano plots displaying differentially expressed genes (DEGs) in Cana fed 12-month-old A) Males; D) Females (p<0.05; FC ≥1.2 and FC≤1/1.2). B) Bubble plot showing KEGG pathway analysis using the DEGs of the Cana fed 12-month-old males. C) GO chord showing DEGs clusters for p53 signaling pathway, MAPK signaling, mTOR signaling, pathways of neurodegeneration, PI3K-AKT signaling, and insulin signaling in Cana fed 12-month-old male mice. E) GO analysis of biological processes, F) GO chord showing cognitive functions and cellular functions related genes, G) Heat map showing gene expression in metabolic and cellular function in the Cana fed 12-month-old female mice. H) Gene ontology (GO) analysis of Biological Processes using the 123 shared genes between the Cana fed 12-month-old males and females. I) GO bubble plot showing the Biological Processes. J) GO chord showing the immune function-related gene expression in 25-month-old male mice. K) GO bubble plot showing the Biological Processes. L) GO chord showing the CNS function-related gene expression in 25-month-old female mice. Shared Cana responsive genes in 12 and 25-month-old M) Male; N) Female mice, n=4/5 animals/group.

### Integrated multi-omics analysis in 12-month-old mice

To get a system-level understanding of Cana’s effects on the hippocampus, we conducted an integrative analysis of the transcriptomics, proteomics, and metabolomics of 12-month-old mice. The Venn diagram illustrates the overlap between significant genes, proteins, and metabolites identified in Cana-treated males (**Figure 5A**). We found an overlap of *Irs4* and *Irs2* genes with elevated IRS1, MGLL, and GNAS1 proteins, alongside the upregulation of the cGMP metabolite, all of which are critical regulators of insulin signaling, metabolic processes, synaptic plasticity, and memory function in the hippocampus (**Figure 5B**). Subsequently, we examined cGMP signaling as a shared mechanistic pathway across omics layers using KEGG annotations (**Figure 5C**). This integrative analysis revealed 52 differentially expressed genes, 28 proteins, and 10 metabolites directly or indirectly associated with cGMP signaling in Cana-treated males. Notably, *Gng3*, *mt-Nd5*, and *Slc24a4* were present in transcriptomic and proteomic datasets. In females, we did not find consistent overlap across the three molecular layers (**Figure 5D and Suppl. File 1)**.

**Figure 5:**
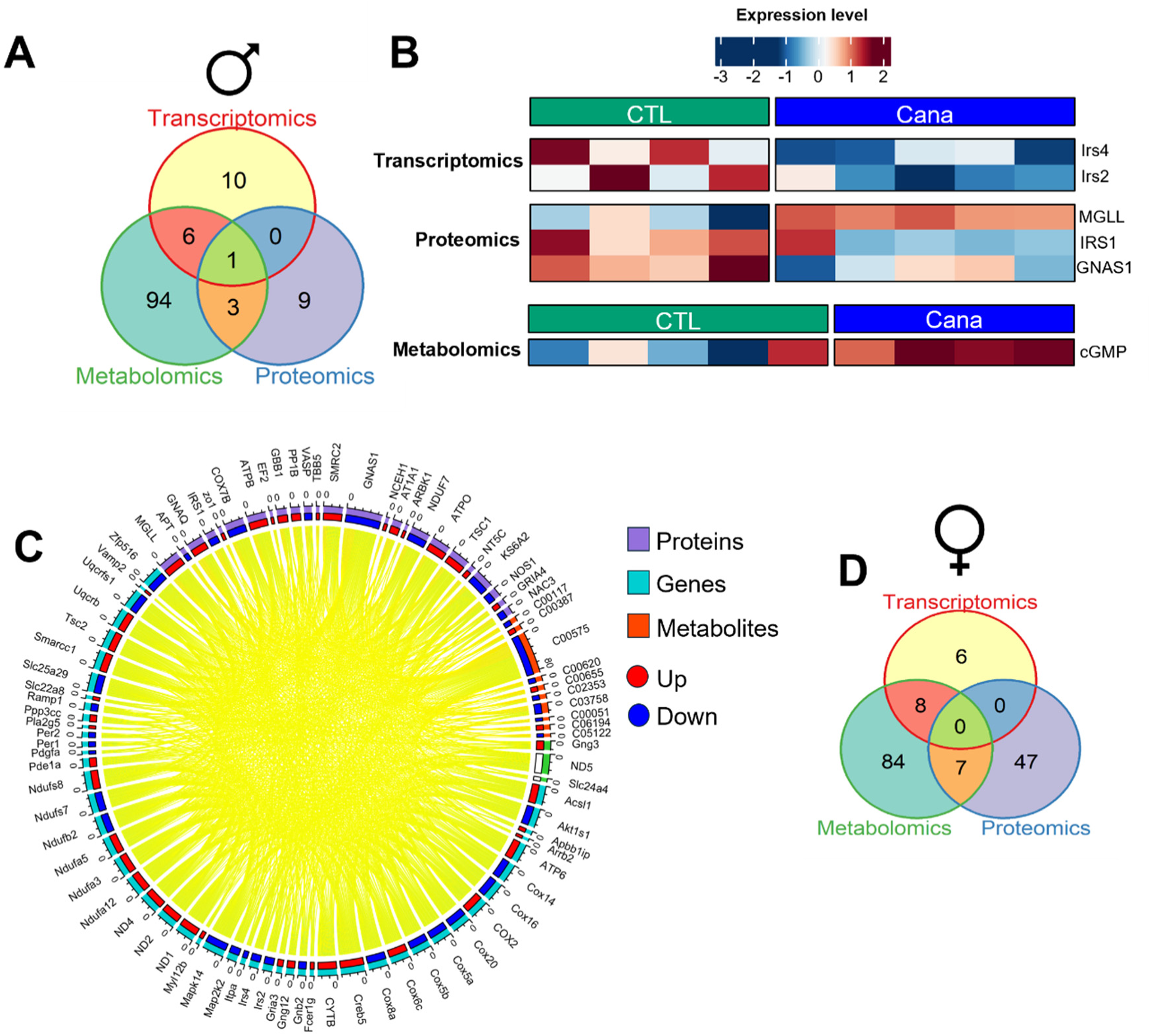
Integrative multi-omics analysis. A) Common and unique significant pathways between transcriptomics, metabolomics, and proteomics data sets in the Cana fed male mice. B) Heat map showing the shared genes, proteins, and metabolites in male mice. C) Chord diagram displaying the relationships among differentially expressed genes, abundant compounds/metabolites, and proteins related to cGMP. D) Ven diagram displaying multi-omics pathways analysis in Cana fed females. n=4/5 animals/group

### Cana treatment delays AD pathology in the 5xFAD mice

The hippocampal response to Cana prompted us to investigate its neuroprotective and metabolic effects in the 5xFAD, a mouse model of AD. Cana treatment started at 3 months of age, before the reported onset of AD pathology in this model(17). The experimental design is illustrated in **Figure 6A**. By 7 months, 5xFAD mice exhibited hyperglycemia, whereas Cana-treated 5xFAD mice of both sexes maintained normoglycemia (**Figure 6B and C**). Body weight, fat mass, and lean mass remained unaffected by genotype, sex, or treatment (**Suppl. Figure 3A-F**). Immunohistochemical analysis showed a significant reduction in Aβ plaque load in the hippocampus of Cana-treated 5xFAD males (effect of Cana p<0.05, sex p<0.05), particularly in the dentate gyrus (DG) and CA3 (effect of Cana p<0.05 and sex p<0.05) subregions. In contrast, the reduction in plaque burden for Cana-treated 5xFAD females didn’t reach significance (**Figure 6D-G**). Analysis of microglial (Iba1^+^) and astrocytic (GFAP^+^) markers revealed sex-specific effects of Cana treatment in 5xFAD mice. In males, Cana significantly reduced microgliosis in both the DG and CA3 sub-regions (effect of Cana p<0.001, and p<0.05, and sex p<0.001), restoring Iba1⁺ cell number to that seen in wild-type controls. CA3 astrogliosis, as indicated by GFAP⁺ staining, was also reduced in Cana-treated males (p<0.01). In contrast, Cana-treated 5xFAD females showed a significant reduction in microglial activation only in the CA3 sub-region (p<0.01), with no change in the DG or GFAP⁺ astrocyte numbers (**Figure 6H-L**). Consistent with this, plaques in 5xFAD males and females were surrounded by activated microglia, positive for Iba1 and CD68, a marker of phagocytic activity (**Suppl. Figure 4**).

**Figure 6:**
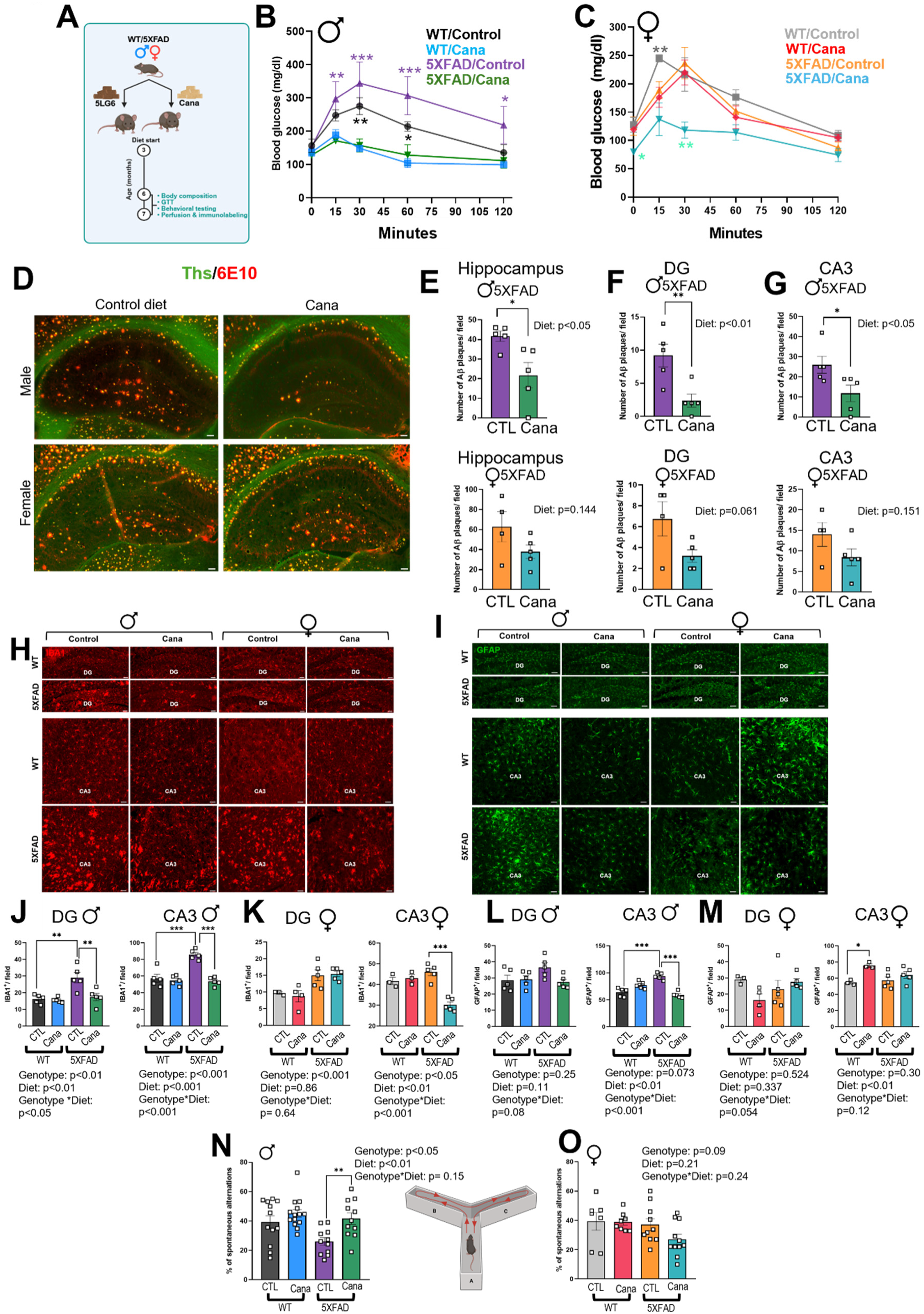
Cana treatment ameliorates Aβ plaque burden, reduces neuroinflammation, and improves spatial memory in 5XFAD male mice. A) 5XFAD experimental design. Glucose tolerance test (GTT) performed on 6-7-month-old B) Male; C) Female mice. D) Representative images demonstrating the Ths (green) and 6E10 (red) positive Aβ plaque distribution in the whole hippocampus and hippocampus sub-regions CA3 and DG in 6-7-months-old mice. Scale bars represent 100 μm. Quantification of the Ths and 6E10 positive Aβ plaques in the E) whole hippocampus and hippocampus sub-regions F) DG, and G) CA3 in 6-7-months-old animals. Representative images showing immunostaining in the CA3, and DG of 6-7-months-old mice for (H) microglia (Iba1^+^ in red) and (I) astrocytes (GFAP^+^ in green). Scale bars: 100 μm. Numbers of cells immunoreactive for Iba1 in the DG and CA3 of (J) Males; (K) Females. Numbers of cells immunoreactive for GFAP in the DG and CA3 of (L) Males; (M) Females of 6-7-month-old WT/5XFAD and Cana/control diet fed animals. n = 3-5 mice/group. Spontaneous alterations were calculated for spatial memory assessment using the Y-maze. (N) Males; (O) Females of 6-7-month-old mice. Error bars show SEM for n = 7-14 mice/group. Data was analyzed using student t-test and Two-factor ANOVA and further analyzed with the Newman–Keuls post hoc test (*p < 0.05, **p<0.01, ***p<0.001).

The Y-maze test on 7-month-old mice assessed working memory. Cana-treated 5xFAD males demonstrated a significantly higher percentage of spontaneous alternation compared to untreated 5xFAD males (p < 0.05), performing similarly to wild-type controls, indicating improved spatial working memory (**Figure 6N**). No differences in spontaneous alternation were found between female groups (**Figure 6O**). At this age, 5xFAD mice showed no deficits in grip strength, rotarod performance, or short-term recognition memory (novel object recognition) (**Suppl. Figure 5**). These findings demonstrate that Cana treatment improved metabolic dysfunction, reduced hallmark AD pathology, and preserved cognitive performance in 5xFAD male mice. Cana’s effects in 5xFAD females were limited to reductions in microgliosis in the CA3 sub-region.

## Discussion

Here, we demonstrate that Canagliflozin (Cana), an FDA-approved SGLT2i, exerts potent neuroprotective effects in aging male mice. Cana treatment enhanced hippocampal-dependent learning and memory, markedly reduced AD pathology, and reprogrammed key molecular pathways linked to energy metabolism, neuroinflammation, and insulin signaling. Using an integrative multi-omics approach, we show integrated activation of neuroprotective and metabolic programs across transcriptomic, proteomic, and metabolomic layers that align with the observed behavioral improvements and neuropathological resilience in aged Cana-treated males. In contrast, females exhibited attenuated molecular responses, with limited functional benefits, demonstrating sex-specific differences in hippocampal aging and drug responsiveness. Our findings indicate that Canagliflozin can serve as a candidate for gerotherapeutic repurposing to preserve cognitive function and delay AD-related pathology during aging.

In UM-HET3 male mice, Cana upregulated key genes and proteins involved in insulin signaling, synaptic function, learning and memory, and the negative regulation of amyloid-beta formation. This molecular profile was accompanied by enhanced cGMP-PKG signaling, glutathione and pyrimidine metabolism, and mitochondrial function, critical pathways for neuronal resilience during aging (35–37). cGMP signaling and pyrimidine metabolism have previously been linked to protection against cognitive decline and AD pathology (38, 39). For example, pharmacological activation of cGMP via phosphodiesterase (PDE) inhibition improves synaptic plasticity and memory in mouse models of AD (40), while enhanced de novo pyrimidine synthesis has been shown to improve synaptic function and protect against cognitive decline in aging and AD animal models(32). Notably, we identified overlapping molecular signatures such as decreased *Irs2* and increased *Gng3*, *Slc24a4*, and *mt-Nd5* expression. Interestingly, *Irs2* (Insulin receptor substrate 2) is a key regulator of downstream insulin signaling, and its deficiency in the brain has been shown to extend lifespan in mice (41). GNG3, a gamma subunit of G-proteins, regulates synaptic transmission and has been linked to age-related cognitive decline and behavioral deficits (42). Notably, *Gng3* knockout mice exhibit susceptibility to seizures, suggesting a role in neuronal excitability and cognitive stability(43). ND5 (NADH dehydrogenase subunit 5), a core component of mitochondrial Complex I, is frequently disrupted in neurodegenerative conditions in human and rodent models (44, 45). The changes in these shared genes across three molecular layers suggest that Cana activates protective programs that may counteract age-related cognitive decline and neurodegeneration.

UM-HET3 Cana-treated males and females exhibited transcriptional remodeling in the hippocampus with aging. However, only males showed clear cognitive and age-related brain improvements in response to Cana treatment. In a previous work, we demonstrated that long-term Cana treatment selectively reduced neuroinflammation and mTOR signaling while enhancing insulin sensitivity in the hippocampus of 25-month-old males but not in females (14). These molecular responses across transcriptomic, proteomic, and metabolomic layers likely underlie the biochemical and behavioral improvements observed in aged Cana-treated males. Interestingly, while Cana-treated females showed transcriptional enrichment in pathways related to insulin signaling, synaptic transmission, and longevity regulation, these changes were primarily driven by the downregulation of key functional genes, including *Akt2*, *Irs3*, *Pdk4*, *Arc*, and *Calb1*. In contrast, at the proteomic level, Cana-treated females exhibited enrichment in pathways such as fatty acid metabolism and ATP synthesis, with upregulation of core proteins in mitochondrial energy production such as ATP5E, ATP5I, MECR, and ASAH1 (46–48). However, this did not translate into cognitive or neuropathological benefit. The dissociation between transcriptomic and proteomic responses in females may reflect a lack of integrated molecular remodeling in neuronal circuits critical for cognitive function.

Proteomic analysis in Cana-treated male mice revealed 40-fold enrichment in the negative regulation of β-amyloid formation, including the upregulation of key proteins such as BIN1, SORL1, RTN3, GGA3, and PIN1. These proteins regulate amyloid precursor protein (APP) trafficking, processing, amyloidogenic cleavage, and clearance, suggesting a coordinated proteostatic response against AD pathology (21–24). Notably, APP and MAPT were also upregulated. However, these proteins do not become pathogenic unless abnormally cleaved (49). This suggests that Cana treatment suppresses early β-amyloid accumulation. Remarkably, we demonstrate that treating 5xFAD mice with Cana before the onset of pathology resulted in a dramatic reduction in hippocampal amyloid-β plaque burden, particularly in the hippocampus, as well as a significant decrease in microgliosis and astrogliosis around the plaques, pathologies typically observed in this model (17). These effects were primarily observed in males suggesting that Cana treatment can suppress early neuroinflammation and amyloid pathology in AD, especially if used early.

Our findings are consistent with previous reports showing that Cana extended lifespan in male but not female mice (10). Interestingly, Cana levels are consistently higher in females than in males’ blood and in the brain, despite matched diet drug levels, including the hippocampus and hypothalamus (10, 14, 50). Regardless, prolonged Cana intake resulted in increased food and water intake and improved energy homeostasis in aged mice of both sexes (50). Higher Cana accumulation in the female hippocampus did not translate into the cognitive or neuropathological benefits observed in males. Long-term Cana treatment induced robust upregulation of genes involved in insulin signaling, neuropeptide signaling, and mitochondrial metabolism in the hypothalamus. These transcriptional changes were more pronounced in males, whereas females exhibited a more attenuated activation of these pathways (50). These differences may reflect sex-specific variation in drug pharmacokinetics and pharmacodynamics (51). It is also possible that higher drug accumulation in females leads to adverse off-target effects, impacting beneficial signaling or inducing metabolic trade-offs over CNS protection (52).

A growing body of clinical evidence suggests that SGLT2i may reduce the risk of dementia in patients with T2D and cardiovascular disease, likely through improvements in glycemic control, cardiovascular function, and inflammation (53). Moreover, recent meta-analyses have reported significantly lower dementia incidence among patients treated with SGLT2i compared to other glucose-lowering drugs (54). While the metabolic benefits of SGLT2i appear similar in both men and women, findings related to dementia risk are more variable. One meta-analysis found a significant reduction in dementia incidence among male patients using SGLT2i, but not in females (55). In contrast, a large population-based study found no significant sex differences, suggesting cognitive protection may occur across sexes (56). However, these findings have been mainly limited to patients with metabolic or cardiovascular diseases, making it unclear whether SGLT2i can have cognitive benefits without metabolic disease (54). Here, we address this gap by demonstrating that Cana improves hippocampal function and molecular resilience in non-diabetic, normally aging mice, demonstrating a neuroprotective role for SGLT2i in brain aging and AD. While several other anti-aging interventions, such as rapamycin and acarbose, have previously shown beneficial effects in AD mouse models (57, 58) only Cana provides translational benefits since it’s already an FDA-approved drug commonly prescribed for T2D with widespread clinical use.

While our findings provide important data on the neuroprotective effects of Cana, several limitations should be acknowledged. We did not dissect the cell-type specific mechanisms underlying Cana’s effects on hippocampal function, such as its effects on the microglia, astrocytes, or neurons. Future studies should incorporate single-cell or spatial transcriptomics to clarify the contributions of each cell type to Cana’s neuroprotective actions. Additionally, the sex-specific responses observed in our study require further investigation. Moreover, while the 5XFAD mouse model provides a valuable tool to assess early AD pathology, it represents a young-onset disease that may not fully capture the slow nature of late-onset AD. Future studies should evaluate the Cana impact in additional AD models. Additional time points and longitudinal analyses will need to be incorporated.

Together, our findings establish Canagliflozin as a promising gerotherapeutic candidate for AD prevention, capable of modulating key neurodegenerative pathways, reducing amyloid pathology, and preserving cognitive function during aging.

## Materials and Methods

This study was designed to evaluate sex-specific effects of Cana on brain aging and AD-like pathology. Both male and female UM-HET3 and 5xFAD mice were included in all experimental groups. All molecular (transcriptomic, proteomic, and metabolomic), behavioral, and histological analyses were conducted separately by sex to identify differential responses to chronic Cana treatment. The inclusion of both sexes enabled the identification of sex-dependent molecular and cognitive outcomes.

### Mouse husbandry

All mice were maintained under standard laboratory conditions with three males or four females per cage from weaning. 5XFAD mice were purchased from the Mutant Mouse Resource and Research Center (MMRRC), supported by NIH (JAX # 03448). 5XFAD hemizygous males were crossed with WT females to obtain the WT and 5XFAD offspring. Mice were provided with food (TestDiet 5LG6: 17.5% protein, 5.6% fat) and water. UM-HET3 mice received a Cana diet at 180 mg/kg of chow from 7 months of age (5LG6 W/10% 180 ppm Cana). 5XFAD and WT animals received the Cana diet from 3 months of age. All mice were provided with water *ad libitum* and housed in temperature-controlled rooms (22°C) on a 12-hour light-dark cycle. Health status checks were conducted regularly in the animal facility.

### Perfusion and immunolabeling

Mice were anesthetized using avertin and perfused with phosphate-buffered saline (PBS) (pH 7.5), followed by 4% paraformaldehyde (PFA). Brains were post-fixed, dehydrated, and then sectioned coronally (30µm) using a sliding microtome, followed by immunofluorescent analysis as described previously(59, 60). For immunohistochemistry brain sections were washed with PBS six times, blocked with 0.3% Triton X-100 and 3% normal donkey serum in PBS for 2h; then the staining was carried out with the following primary antibodies overnight: mouse anti-6E10 (1:200; Biolegend Cat. No. SIG-39320; rabbit anti-Iba1 (1:1000 Abcam Cat. No. ab5076); goat anti-GFAP (1:1000 Sigma-Aldrich Cat. No. AB5804). For rabbit anti-CD68 (1:300 Abcam Cat No. ab125212) together with Iba1 (1:1000 Abcam Cat. No. ab5076) immunostaining sections were pretreated with 0.5% NaOH and 0.5%H2O2 in PBS for 20 min followed by the glycine treatment for 10 min and blocking with 0.3% Triton X-100 and 3% normal donkey serum in PBS for 1h. Following the primary antibody treatment brain sections were incubated with AlexaFluor-conjugated secondary antibodies for 2h. For the Aβ plaque staining sections were incubated for 5 mins in Ths (500 µg; Sigma-Adrich) dissolved in 50% ethanol after the secondary antibody incubation and washing. Microscope images of the stained sections were obtained using Nikon 800 fluorescent microscope using Nikon imaging DS-R12 color-cooled SCMOS, version 5.00 and Laser scanning confocal microscope Zeiss LSM 800.

### Quantification

For the cell quantification and immunoreactivity analysis, images were taken from at least 4 sections containing the hippocampus for each brain between bregma −0.82 mm to −2.4 mm (according to the Franklin mouse brain atlas). Serial brain sections were made at a 30 µm thickness. Fiji-ImageJ was used to count 6E10 and Ths colocalized Aβ plaques and GFAP and Iba1 positive cells. For phagocytotic microglia visualization, confocal images of Ths, CD68, and Iba1 were taken using the multiphoton laser-scanning microscope (LSM 800, ZEISS) with 40X and 63X objectives.

### Behavioral assays

All the behavioral testing was conducted with a two-day rest period between each test. The tests were conducted in the following order: open field test, spontaneous alternation, rotarod, grip strength, accelerated rotarod, novel object recognition test and Barnes maze. On each test day, animals were transported from the housing room to the procedure room, and the animals were left to acclimate to the procedure room and testing environment for at least 1 hour. Animals were provided with ad libitum water and food in the procedure room and the housing room. Silence in the room was maintained throughout the testing time to avoid any disturbance or stress to the animal’s natural behavior. Between each test arena, the area was sanitized with 70 % ethanol and dried before introducing the next animal to the arena. Lighting in the testing room was consistent with the housing room except for the novel object test, which was conducted under ambient lighting (∼20-35 lux). Animals’ movement within the arenas was recorded using 22 Series CMOS camera and the recorded videos were analyzed using the Any-Maze video tracking system^©^ V 7.49. A brief description of each behavioral test is provided below.

### Open field maze

The open field arena was 31.7”×31.7”×11.6” and made with polyvinyl chloride (PVC). The center of the arena was defined as a 16”×16” section from the middle and the four corners of the arena were defined as the corners of the arena. The animals inside the arena were recorded for 10 minutes. As the test started animals were placed in the center of the arena and allowed to walk freely. After each test, the animals were returned to their home cage. The travel distance and time spent in the center for each animal were calculated using the Any-maze software.

### Spontaneous Alternation

The Y-maze arena was 13.5”×2”×7.8” and made with polyvinyl chloride (PVC). The three arms were labeled as “A”, “B”, and “C” for later analysis purposes. Animals were placed halfway in the start arm (A) facing the center of the Y, and allowed to explore the Y-maze for 8 min. The sequence of the entries into each arm was recorded. Using the Any-maze video tracking software, the % of spontaneous alterations was calculated using the equation below.

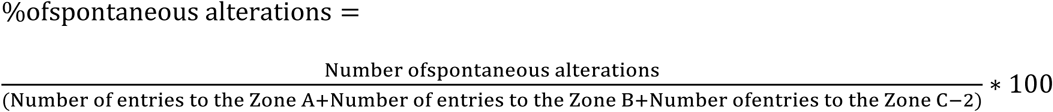

### Barnes maze

Barnes maze arena was 18” radius with 20 holes around arena’s edge and four distinct visual cues. The test was conducted in three phases: acquisition phase, short-term memory (STM) test and long-term memory (LTM) test. On the first day before the acquisition starts animals were introduced to the arena, guided to the escape hole, and allowed to stay inside the escape hole for 2 min. Then, three acquisition trials were conducted for each day of the acquisition phase, with a maximum time of 180 seconds for each trial. During the acquisition phase, the location of the escape hole and visual cues weren’t changed. On the 5^th^ day, the STM test was performed, and on the 12th day, the LTM test was performed. During the STM and LTM, the escape hole was removed, and the animal was introduced to the middle of the arena and allowed to find the escape hole location within 90 seconds. All these phases were conducted while a white noise was in the background. Latency to target hole and time spent at the target hole were analyzed using the Any-maze video tracking software.

### Hippocampus RNA extraction

Male mice were sacrificed at ad libitum to harvest the brain and to isolate the hippocampus. Hippocampus samples were lysed with 0.75 ml of 2-mercaptoethanol-supplemented lysis buffer (PureLink® RNA Mini Kit #12183025). After the homogenization, with 0.75 ml of 70% ethanol, samples were transferred to spin cartridges with the collection tubes. Samples were centrifuged at 12000 × g for 15 s at room temperature. After washing the samples two times with washing buffers, the cartridges were centrifuged at 12000 × g for 1–2 min to dry the membrane with bound RNA. RNA was eluted using 15–20 μL of RNase-free water.

### RNA sequencing and data analysis

Hippocampus RNA-Seq was performed at the WSU Genome Sciences Core. RNA concentration was determined by NanoDrop (ND-1000 UV–Vis Spectrophotometer) and quality was assessed using RNA ScreenTape on a 4200 TapeStation. RNA-seq libraries were prepared according to the Illumina Stranded Total RNA Prep with Ribo-Zero Plus protocol and sequenced on a NovaSeq 6000 (50 bp x2) to an average depth of 30M raw paired reads per transcriptome. Adapter trimming and low-quality sequences were removed from the reads using Trim Galore v0.6.0 with the ‘--paired’ parameter. Processed reads were aligned to the mm10 genome with GENCODE M24 gene annotations using STAR v2.7.5a default parameters to generate read counts per gene.

Reads were tabulated to gene regions using HTSeq-0.6.1p1. Males and females were analyzed separately. RNA-seq data were processed using the edgeR package (61) in the statistical software R. Missing values and normalization were handled using the trimmed mean of M-values (TMM) method implemented in edgeR. Proteomics data were preprocessed using random forest imputation(62) for missing values, followed by median-based sample normalization. Expression and abundance values from transcriptomic and proteomic datasets were log2-transformed before analysis. Principal component analysis (PCA) assessed the separation between experimental groups. Two- and three-dimensional PCA plots were generated using the first two or three principal components. Differential expression and abundance were assessed and visualized using volcano plots, applying an unadjusted p-value threshold of 0.05 and a fold-change cutoff of 1.2. Significantly differentially expressed genes (DEGs) were generated using the R Studio package DESeq2. Statistical significance was calculated by adjusting the P values with the Benjamini-Hochberg’s false discovery rate (FDR) method. DEGs were used to identify the enriched pathways, both Gene Ontology (for Biological Processes (BP)) and KEGG enrichment pathways using Gene Set Enrichment Analysis (GSEA) (NIH DAVID Bioinformatics https://david.ncifcrf.gov/home.jsp) (p-value cut off < 0.05). All RNA-Seq data are available at the Sequence Read Archive (SRA) at NCBI under accession number PRJNA1263876.

### Hippocampus protein extraction

Hippocampus tissue was homogenized using a Stom 24 Bullet Blender (Next Advance) with 100 µL of unbuffered water and 3.2 nm stainless steel beads. After the initial homogenization step, samples were again homogenized with 100 µL of 5% lithium dodecyl sulfate (LiDS) detergent. Then the homogenate was heated at 95 °C for 5 minutes and then filtered through spin columns (Pierce #89868). A small aliquot from each clear filtered lysate was used for protein quantification. The remaining lysate was buffered with 20 mM tetraethylammonium bicarbonate (TEAB; Honey well Fluka Cat. No. 60-044-974), reduced with 5 mM dithiothreitol (DTT), and alkylated with 15 mM iodoacetamide (IAA). Excess IAA was quenched after 30 min by adding another aliquot of 5 mM DTT. Samples were acidified by adding 20 µL of 12% phosphoric acid, followed by protein precipitation using 1 mL of 90% methanol (MeOH) in 100 mM TEAB. The resulting protein pellets were washed with 0.5 mL of 80% MeOH in 10 mM TEAB, air-dried at room temperature, and subsequently resuspended in 150 µL of buffer containing 100 mM NaCl, 1 mM CaCl₂, 40 mM TEAB, and 0.5% deoxycholate (DOC). Finally, proteins were digested by adding 1 µg of trypsin (Promega, V5113) per sample, followed by overnight incubation at 37°C.

### Mass spectrometry and proteomic data analysis

LC-MS/MS analysis was conducted using a Thermo Scientific Vanquish-Neo chromatography system equipped with an Acclaim PepMap 100 trap column (100 µm × 2 cm, C18, 5 µm, 100 Å) and an Easy-Spray PepMap RSLC C18 analytical column (75 µm × 25 cm). All samples were analyzed using a gradient starting from 6% to 42% acetonitrile (80% solution in 0.1% formic acid) for 60 minutes. Mass spectrometry was performed on an Orbitrap Eclipse system utilizing data-independent acquisition (DIA). MS1 spectra were acquired with a mass range of 350–1650 Da at 120,000 resolution, using an AGC target of 3e6. MS2 spectra, obtained through higher-energy collisional dissociation (HCD) fragmentation at a collision energy of 30, covered a 200–1600 Da range at 15,000 resolution, with a maximum injection time of 120 ms and an AGC target of 3e6. Spectral data were processed using Spectronaut 18.6 (Biognosys) with default BGS factory settings normalization, referencing the Mouse UniProt FASTA database (downloaded March 30, 2021; 17,035 entries). Trypsin was used as the digestion enzyme, allowing up to two missed cleavages. Carbamidomethylation of cysteine was designated as a fixed modification, while oxidation (M), acetylation (protein N-terminus), methionine loss, or both acetylation and methionine loss were included as variable modifications. A false discovery rate (FDR) cutoff of 1% was set for precursor and peptide identification. Proteins exhibiting over 50% missing values across the 19-sample dataset were excluded, and remaining missing values were imputed using a random forest algorithm. Protein abundance values were normalized to the median intensity of each sample and log2-transformed to ensure variance stabilization. Quantitative differences among sample groups were subsequently assessed by Principal Component Analysis (PCA) and Partial Least Squares-Discriminant Analysis (PLS-DA).

### Hippocampus metabolite extraction

Dissected hippocampus samples stored at −80 °C were thawed on ice and homogenized in a ball-mill grinder at 30 Hz for 20s. 400 µL solution (Methanol: Water = 7:3, V/V) containing internal standard was mixed with 20 mg of ground sample and mixed in a shaker at 2500 rpm for 5 min. The mixture was then placed on ice for 15 min and centrifuged at 12000 rpm for 10 min (4 °C). Then, 300 µL of the supernatant was collected and placed in −20 °C for 30 min. The samples were centrifuged at 12000 rpm for 3 min (4 °C). A 200 µL aliquot of the supernatant was used for LC-MS analysis.

### Mass spectrometry and metabolomic data analysis

The original data file acquired by LC-MS was converted to mzML format by ProteoWizard. The XCMS program performed peak extraction, peak alignment, and retention time correction. The peaks with a missing rate >50% in each group of samples were filtered. The blank values were filled with KNN, and the peak area was corrected using the SVR method. The metabolites were annotated by searching MetwareBio’s in-house database, integrated public, prediction, and metDNA databases. Finally, substances with a comprehensive identification score above 0.5 and a CV value of QC samples less than 0.3 were extracted. Then, positive and negative modes were combined (substances with the highest qualitative grade and the lowest CV value were retained). Based on the results of OPLS-DA (biological repetition ≥ 3), multivariate analysis of Variable Importance in Projection (VIP) from OPLS-DA modeling was used to select differential metabolites from different samples. Differential metabolites from univariate analysis can be screened by combining the P-value/FDR (when biological replicates ≥ 2) or FC values. The screening criteria for this project are as follows: Metabolites with VIP > 1 were selected. VIP value represents the effect of the differences between groups for a particular metabolite in various models and sample groups. It is generally considered that the metabolites with VIP > 1 are significantly different, and Metabolites with P-value < 0.05 (Student’s t test was used when the data follows a normal distribution, otherwise Wilcoxon rank-sum test) were considered as significant differences and selected.

### Statistical analysis

Data sets were analyzed using Student’s t-test and two-way analysis of variance (ANOVA), followed by Newman-Keuls post hoc test. All data were presented as mean ± SEM. P < 0.05 was considered significant. TIBCO Statistica^®^ v. 13.5.0.17 was used for statistical analysis.

### Study Approval

All animal experiments followed NIH guidelines for Animal Care and Use, approved and overseen by Wayne State University Institutional Animal Care and Use Committee (IACUC).

## Author contribution

HJ, DHM, NC, OK carried out the research and analyzed the data. KG, SK, and HJ analyzed the genomic, metabolomic, and proteomic data. BCG and SY analyzed the data, reviewed, and revised the manuscript. RAM provided the UM-HET3 mice and Cana diet, reviewed, and revised the manuscript. MS designed the study, analyzed the data, wrote the manuscript, and is responsible for the integrity of this work. All authors approved the final version of the manuscript.

## Data Availability Statement

The data that support the findings of this study are available from the corresponding author upon reasonable request.

## Abbreviations

Cana: Canagliflozin
CNS: central nervous system
SGLT2i: sodium-glucose co-transporter 2 inhibitor
AD: Alzheimer’s disease
APP: amyloid precursor protein
STM: short-term memory
LTM: long-term memory
DG: dentate gyrus.

