## Supplementary Figures for "Canagliflozin reprograms the aging hippocampus in genetically diverse UM-HET3 mice and attenuates Alzheimer’s-like pathology"

**Running title:** Canagliflozin effect on aging hippocampus

**Keywords:** Canagliflozin, aging brain, hippocampus, metabolism, Alzheimer's disease, longevity

**Corresponding author:**

***Marianna Sadagurski,**
**Department of Biological Sciences,**
**Integrative Biosciences Center**
**Wayne State University**
**6135 Woodward, Detroit, MI 48202**


#### Supplementary Material and Methods

##### Grip strength and Rotarod tests

Animals were tested for forepaw grip strength using DS2-50N grip strength meter. The animals were removed from their home cages by the base of their tail and suspended above the grid until their forepaws gripped the grid. Then the animals were pulled down from their tail gently, horizontally away from the grid until the animal released its grip. The maximum force was recorded. Each animal was tested five times with a 10-second rest between each. The average of five tests was used for the analysis. Animals were also tested for their ability to balance on an accelerating rotarod. The protocol was set for the rotarod to gradually increase its speed to reach 40 revolutions per minute (RPM) within 5 minutes. Then the animals were removed from their home cages and introduced to the rotarod. At the beginning of the test all the animals were facing the forward side of the rotarod. The time and speed at which animals dropped from the accelerating rotarod were used as scores; each animal was tested three times, with 5 min intervals between trials. average of the tests was used for the analysis.

##### Novel object recognition (NOR)

NOR arena was 15.2”×15.2”×11.6”. Novel object was conducted over 3 days with a habituation familiarization and novel object phase. During the first two days animals were transported to the procedure room and allowed to acclimatize for 1 hour. Then the animals were introduced into the empty arena for 10 minutes to become habituated to the arena. The habitation was done for two days. On the third day, two identical objects were placed in the arena diagonally, and the animals were introduced between the identical objects. The animals were allowed to explore identical objects for 10 minutes. After the 10 minutes of familiarization animals were put back in their cages and left undisturbed for 3 hours. After 3 hours, one of the identical objects was switched to a novel object, and the animals were again introduced to the arena between the objects. Animals were again allowed to explore the objects for 10 minutes. During identical and novel object exploration, animals were recorded and the time spent at the novel object and the discrimination index (below equation) were analyzed using Any-maze software.

$$\text{Discrimination index} = \frac{(\text{Time investigating the novel object zone} - \text{Time investigating the familiar object zone}}{(\text{Time investigating the novel object zone} + \text{Time investigating the familiar object zone}}$$

**Measurement of Canagliflozin in mouse hippocampus using HPLC/MS/MS**

All the measurements were performed as previously described[1,2]. The HPLC/MS/MS system consisted of a Shimadzu SIL 20A HT autosampler, LC-20AD pumps and an AB Sciex API 4000 tandem mass spectrometer, with turbo ion spray. The LC analytical column was an ACE Excel C18-PFP (75 x 3.0 mm, 3 microns) purchased from Mac-Mod Analytical (Chaddsford, PA) and was maintained at 25 °C during the chromatographic runs using a Shimadzu CT-20A column oven. Mobile phase A contained 0.1% formic acid dissolved in water. Mobile phase B contained 0.1% formic acid dissolved in 100% HPLC grade acetonitrile. The flow rate of the mobile phase was 0.4 ml/min. Canagliflozin and Lidocaine D10 (an internal standard obtained from Sigma Aldrich, St. Lous, MO) were eluted with a gradient. The initial mobile phase was 20% B and at 1 minute after injection was ramped to 100% B. From 4.0 min to 8.0 min the mobile phase was maintained at 100% B and 8.01 minutes was switched immediately back to 20% B and ran for 1.99 minutes to equilibrate the column before the next injection. The Cana transition was detected in positive mode at 445.3/267.2 Da. The internal standard, Lidocaine D10, transition was detected at 245/96 Da. Each hippocampal section (39 ± 7 mg, mean ± SD) was weighed in a polypropylene tube and then a 10x volume of 75% methanol was added. Each sample was thoroughly homogenized. Calibrator samples were prepared by spiking a control brain homogenate to achieve final concentrations of 0, 10, 20, 40, 80, 100, and 400 ng/ml. Briefly, 0.2 mL of the calibrator and unknown brain homogenates were mixed with 10 µL of 1 µg/mL lidocaine D10. The samples were vortexed vigorously and then centrifuged at 13,000 g for 5 min at 25 °C. The supernatants were transferred to clean 1.5 mL microcentrifuge tubes and centrifuged at 13,000 g for 2 min at 25 °C. The samples were transferred to injection vials, and 10 µL was injected into the HPLC/MS/MS. The concentration of Cana was expressed as ng/mg brain.

#### Supplementary Legends

**Supplementary Figure 1: Principal component analysis (PCA) of the hippocampus.** Proteomics (A) Males, (B) Females. Metabolomics (C) Males, (D) Females. Transcriptomics (E) Males, (F) Females of 12 months of age, (G) Males and (H) Females of 25 months of age.

**Supplementary Figure 2: Hippocampal transcriptomic analysis in Cana-fed mice.** (A) GO chord showing the differentially expressed genes (DEGs) in KEGG pathways in 12-month-old males. B) GO chord showing the differentially expressed genes (DEGs) in nervous system function of a 25-month-old male. Heat map showing genes involved in metabolism and cellular signaling in (C) Males; (D) Females of 25 months of age.

**Supplementary Figure 3: Body composition analysis of 5XFAD mice.** Body weights of (A) Males and (D) Females; Fat mass of (B) Males and (E) Females; Lean mass of (C) Males and (F) Females of 6-7-month-old WT/5XFAD and Cana/control fed mice. Error bars show SEM for n = 5-10 mice/group.

**Supplementary Figure 4: Activated microglia, positive for CD68, a marker of phagocytic activity in 5XFAD mice.** Representative confocal image of 5xFAD Cana or control-fed male and female mice stained with Ths (green), Iba1 (red), and Cd68 (purple).

**Supplementary Figure 5: Behavioral assessment of 5XFAD mice.** All the tests were conducted on 6-7-month-old WT/5XFAD and Cana/control-fed mice. Travel distance (A) Males; (B) Females; Time spent in the center (C) Males; (D) Females during the open field test. Latency to drop (E) Males; (F) Females. Drop speed (G) Males; (H) Females during the rotarod test. Grip strength (I) Males; (J) Females. Discrimination index of (K) Males; (L) Females during the novel object recognition test. Error bars show SEM for n = 7-14 mice/group.

1 Jayarathne, H. S. M. *et al.* Neuroprotective effects of Canagliflozin: Lessons from aged genetically diverse UM-HET3 mice. *Aging Cell* **21**, e13653 (2022). https://doi.org/10.1111/acel.13653
2 Mohamed, D., Elshahed, M. S., Nasr, T., Aboutaleb, N. & Zakaria, O. Novel LC-MS/MS method for analysis of metformin and canagliflozin in human plasma: application to a pharmacokinetic study. *BMC Chem* **13**, 82 (2019). https://doi.org/10.1186/s13065-019-0597-4

Suppl 1

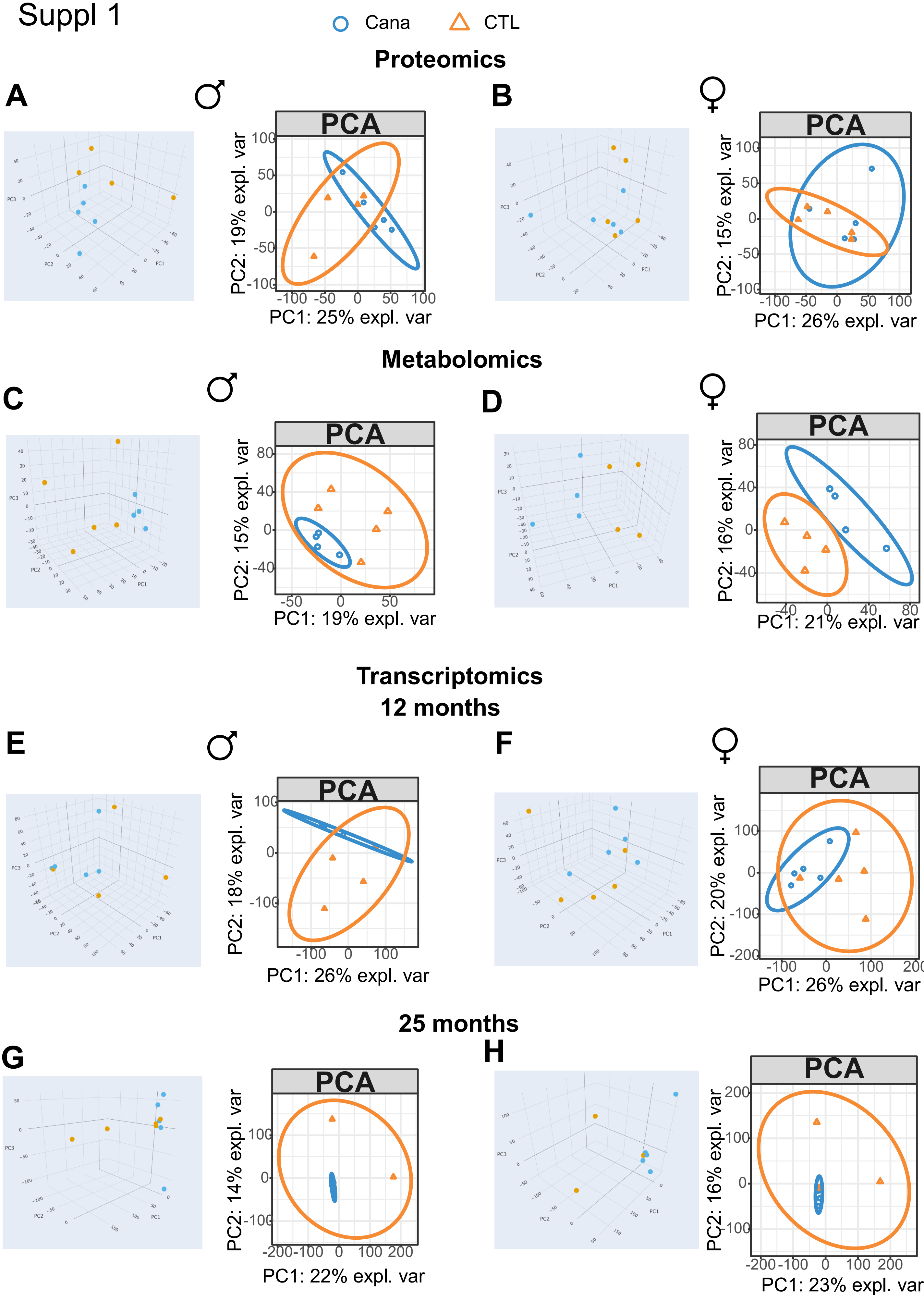

Cana
CTL
Proteomics
A
B
PCA
PC2: 19% expl. var
PC1: 25% expl. var
PC2: 15% expl. var
PC1: 26% expl. var
Metabolomics
C
D
PC2: 15% expl. var
PC1: 19% expl. var
PC2: 16% expl. var
PC1: 21% expl. var
Transcriptomics
12 months
E
F
PC2: 18% expl. var
PC1: 26% expl. var
PC2: 20% expl. var
PC1: 26% expl. var
25 months
G
H
PC2: 14% expl. var
PC1: 22% expl. var
PC2: 16% expl. var
PC1: 23% expl. var

### Suppl 2

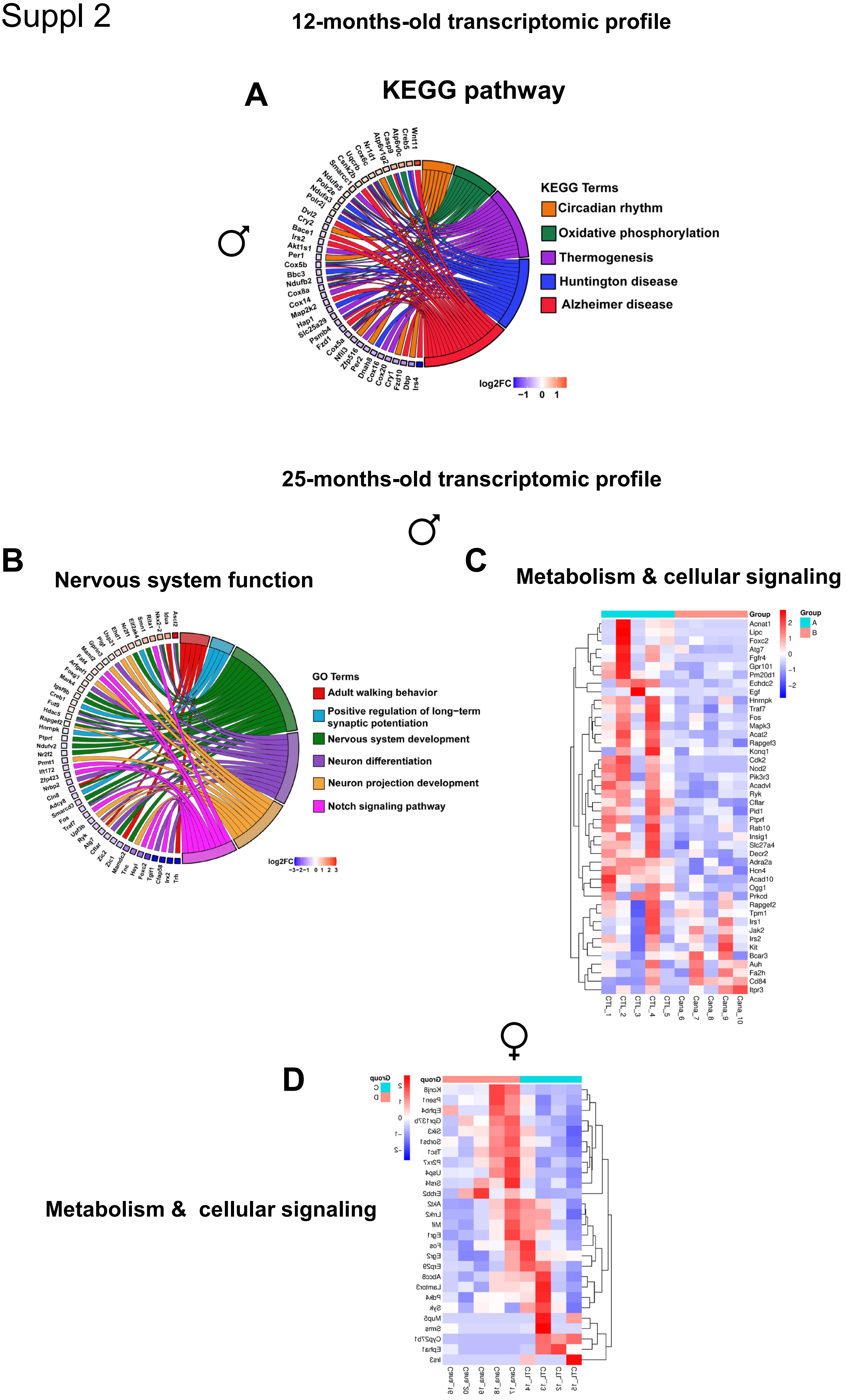
12-months-old transcriptomic profile
A
KEGG pathway
♂
KEGG Terms
Circadian rhythm
Oxidative phosphorylation
Thermogenesis
Huntington disease
Alzheimer disease
log2FC
−1 0 1
Wnt11
Creb5
Atp6v0c
Casp9
Atp6v1g2
Nr1d1
Cox6c
Uqcrb
Csnk2b
Smarcc1
Ndufa5
Polr2e
Ndufa3
Polr2j
Dvl2
Cry2
Bace1
Irs2
Akt1s1
Per1
Cox5b
Bbc3
Ndufb2
Cox8a
Cox14
Map2k2
Hap1
Slc25a29
Psmb4
Fzd1
Cox5a
Nfil3
Zfp516
Per2
Dnah8
Cox16
Cox20
Cry1
Fzd10
Dbp
Irs4
25-months-old transcriptomic profile
♂
B
Nervous system function
GO Terms
Adult walking behavior
Positive regulation of long−term synaptic potentiation
Nervous system development
Neuron differentiation
Neuron projection development
Notch signaling pathway
log2FC
−3−2−1 0 1 2 3
C
Metabolism & cellular signaling
Group
A
B
Acnat1
Lipc
Foxc2
Atg7
Fgfr4
Gpr101
Pm20d1
Echdc2
Egf
Hnrnpk
Traf7
Fos
Mapk3
Acat2
Rapgef3
Kcnq1
Cdk2
Nod2
Pik3r3
Acadvl
Ryk
Cflar
Pid1
Ptprf
Rab10
Insig1
Slc27a4
Decr2
Adra2a
Hcn4
Acad10
Ogg1
Prkcd
Rapgef2
Tpm1
Irs1
Jak2
Irs2
Kit
Bcar3
Auh
Fa2h
Cd84
Itpr3
CTL_1
CTL_2
CTL_3
CTL_4
CTL_5
Cana_6
Cana_7
Cana_8
Cana_9
Cana_10
♀
D
Metabolism & cellular signaling

Suppl 3

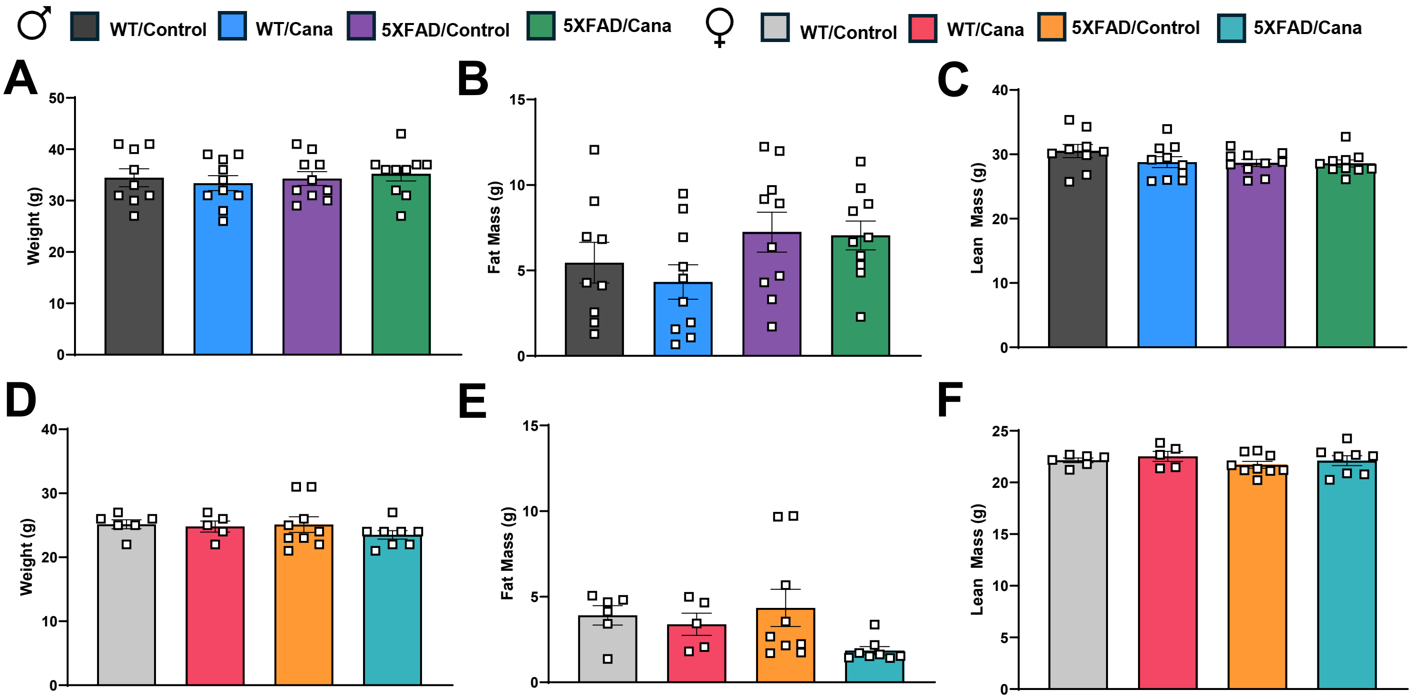

♂
WT/Control
WT/Cana
5XFAD/Control
5XFAD/Cana
♀
WT/Control
WT/Cana
5XFAD/Control
5XFAD/Cana
A
Weight (g)
0
10
20
30
40
50
B
Fat Mass (g)
0
5
10
15
C
Lean Mass (g)
0
10
20
30
40
D
Weight (g)
0
10
20
30
40
E
Fat Mass (g)
0
5
10
15
F
Lean Mass (g)
0
5
10
15
20
25

Suppl 4

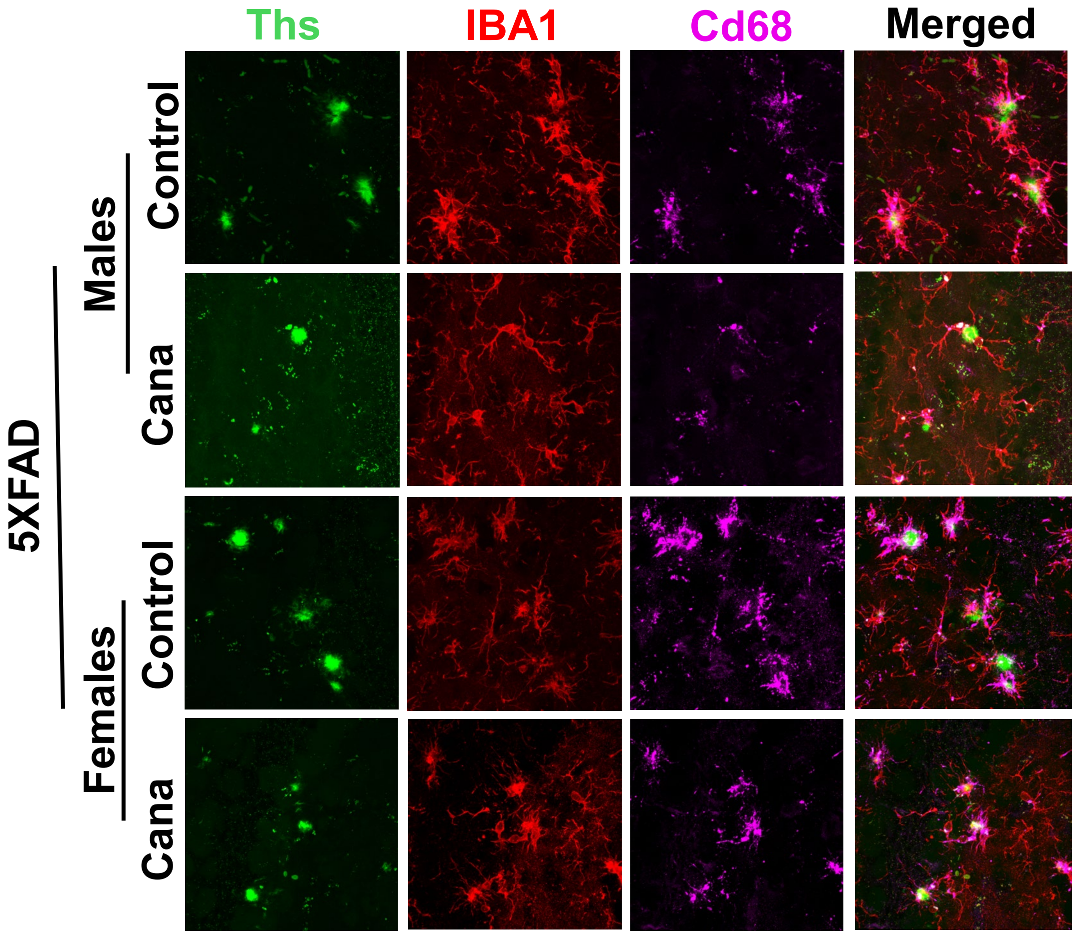

Ths
IBA1
Cd68
Merged
5XFAD
Males
Control
Cana
Females
Control
Cana

Suppl 5

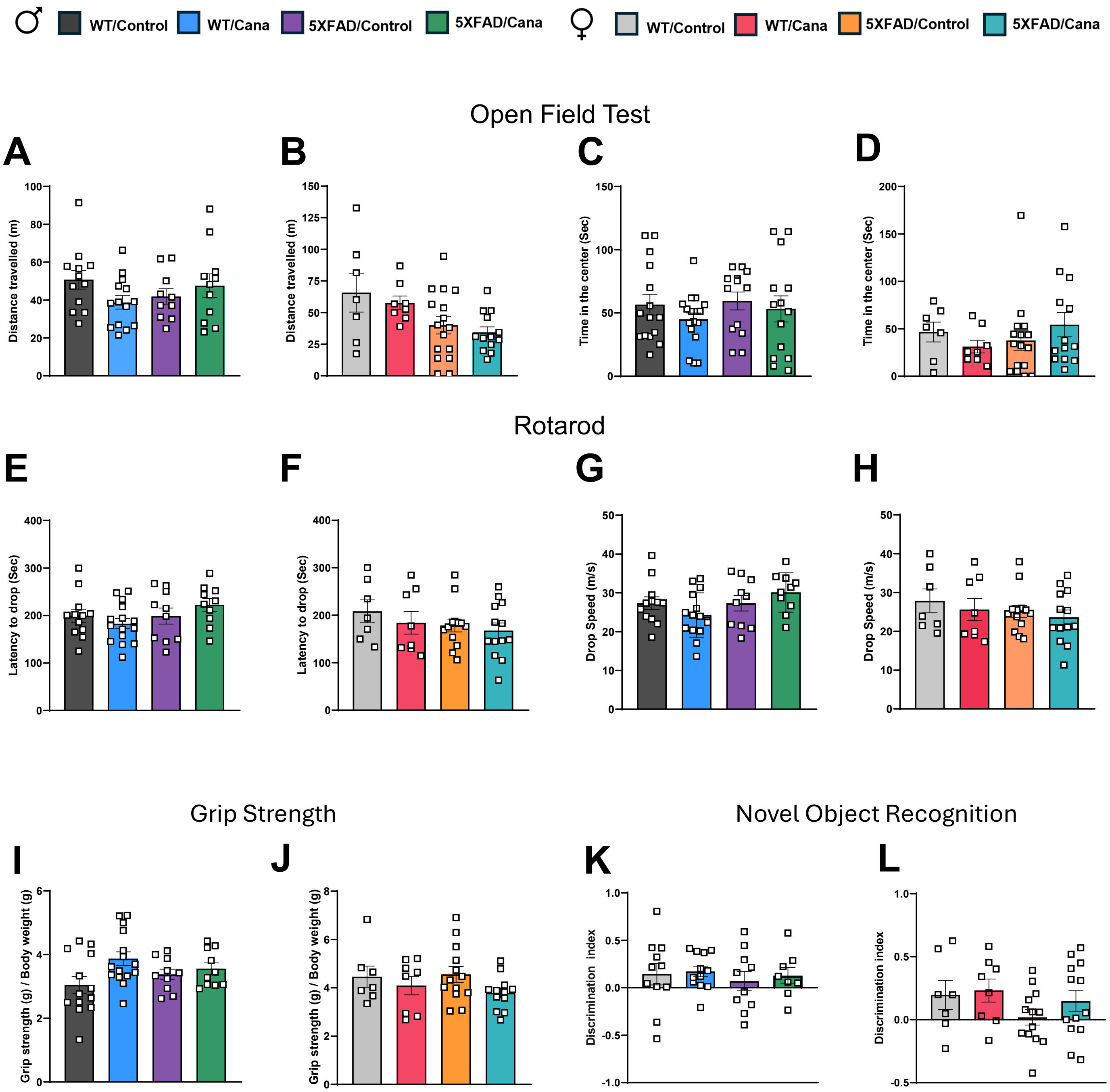

♂
WT/Control
WT/Cana
5XFAD/Control
5XFAD/Cana
♀
WT/Control
WT/Cana
5XFAD/Control
5XFAD/Cana
Open Field Test
A
Distance travelled (m)
B
Distance travelled (m)
C
Time in the center (Sec)
D
Time in the center (Sec)
Rotarod
E
Latency to drop (Sec)
F
Latency to drop (Sec)
G
Drop Speed (m/s)
H
Drop Speed (m/s)
Grip Strength
I
Grip strength (g) / Body weight (g)
J
Grip strength (g) / Body weight (g)
Novel Object Recognition
K
Discrimination index
L
Discrimination index
